# Cross-kingdom miRNA delivery by Panax notoginseng-derived EVs restores neuronal function after ischemic injury

**DOI:** 10.1101/2025.08.08.669445

**Authors:** Yuanyuan Yu, Na Tan, Zhifeng Xu, Zhijian Tan, Tao Wang, Huimin Liu, Le Xu, Dan Lu, Yamei Tang, Hongcheng Mai

**Author notes:** Co-corresponding authors: Dan Lu; Yamei Tang; Hongcheng Mai. These authors contributed equally to this work (Yuanyuan Yu, Na Tan, Zhifeng Xu).

## Abstract

The impermeability of the blood–brain barrier (BBB) remains a major obstacle to effective treatment of neurological disorders, particularly ischemic stroke. Here, we show that plant-derived extracellular vesicles (PEVs) offer a promising strategy to overcome this barrier. Using an optimized high-yield extraction protocol, we isolated PEVs from four medicinal plants—*Panax ginseng, Panax notoginseng, Gastrodia elata,* and *Ligusticum chuanxiong.* Among these, EVs derived from *Panax notoginseng* (NotoEVs) exhibited the strongest neuroprotective effects under hypoxic conditions in both in vitro and in vivo stroke models. Mechanistically, NotoEVs deliver conserved plant microRNAs to recipient neurons, where they suppress key stress granule nucleators (G3bp2, Ubap2l, Lsm14a), activate mTOR signaling, and promote mitochondrial stabilization via the BCL2/TOM20 axis. This cross-kingdom RNA delivery reprograms neuronal stress responses, reduces infarct volume, preserves neuronal morphology, and restores electrophysiological function. Together, our findings establish a scalable platform for plant-based nanotherapeutics and highlight the translational potential of NotoEVs for the treatment of ischemic stroke.

## Introduction

Ischemic stroke (IS) remains a major cause of death and disability worldwide [1], with 69.9 million cases reported in 2021 and an age-standardized incidence rate of 92.4 per 100,000 individuals [2]. Despite advances in acute interventions such as thrombolysis, therapeutic options are still limited by a narrow treatment window [3–4] and the risk of hemorrhagic transformation [5–6]. The limited permeability of the blood-brain barrier (BBB) further complicates effective drug delivery, restricting the utility of many promising neuroprotective agents [7–8].

Plant-derived extracellular vesicles (PEVs) have recently emerged as potential therapeutic agents for overcoming these barriers [9]. Nanoscale vesicles (30–150 nm in diameter) are naturally enriched with bioactive molecules, including proteins and RNAs, and have demonstrated low immunogenicity and intrinsic tissue-targeting capabilities. Initial studies have suggested that PEVs may confer neuroprotective effects in cerebrovascular and neurodegenerative diseases [10–11]; however, low extraction yields and insufficient standardization have hindered their clinical translation.

To address these limitations, we developed an optimized extraction strategy combining differential ultracentrifugation with sucrose density gradient purification, achieving yields exceeding 10^12^ particles/mL from fresh rhizome tissues. This advancement has enabled the isolation and functional characterization of a high-purity PEV population derived from *Panax notoginseng* (NotoEVs).

In this study, we investigated the ability of NotoEVs to promote neuronal recovery after ischemic injury. Using a combination of oxygen-glucose deprivation/reperfusion (OGD/R) and *in vivo* stroke models, we demonstrated that NotoEVs preserved neuronal morphology, restored mitochondrial function, and enhanced synaptic plasticity. Mechanistically, we identified a critical role for plant-derived microRNAs (miRNAs) in modulating stress granule dynamics and activating mTOR-dependent pathways, revealing a previously unrecognized cross-kingdom regulatory mechanism underlying neuroprotection. This insight into PEV-associated miRNAs offers a new understanding of how plant vesicles confer resilience to neural cells, opening exciting possibilities for plant-based nanotherapeutics for stroke and other neurological disorders.

## Results

### Efficient isolation and characterization of PEVs

In this study, by optimizing the extraction process, PEVs were successfully isolated from the fresh rhizome tissues of *Panax ginseng*, *Panax notoginseng*, *Gastrodia elata*, and *Ligusticum chuanxiong* (Figure 1A). Conventional exosome isolation methods face significant limitations: ultrafiltration is time-consuming, cost-ineffective, and causes substantial sample loss — particularly unsuitable for viscous samples — while ultracentrifugation proves cumbersome for large-volume specimens. To overcome these challenges, we developed an innovative integrated ultrafiltration-ultracentrifugation-sucrose cushion (UF-UC-SC) approach comprising three sequential steps: (1) volume reduction via ultrafiltration, (2) exosome-enriched pellet collection by ultracentrifugation (containing exosomes with minor impurities), and (3) purification through sucrose cushion centrifugation after resuspension in minimal-volume PBS. This methodology reduced ultracentrifugation cycles by 50%, shortened processing time by 40%, and decreased sample loss by >60% compared to classical ultracentrifugation. Critically, exosomes isolated via this approach maintained structural integrity, as confirmed by transmission electron microscopy revealing characteristic cup-shaped morphologies with >90% vesicle integrity.

**Figure 1.**
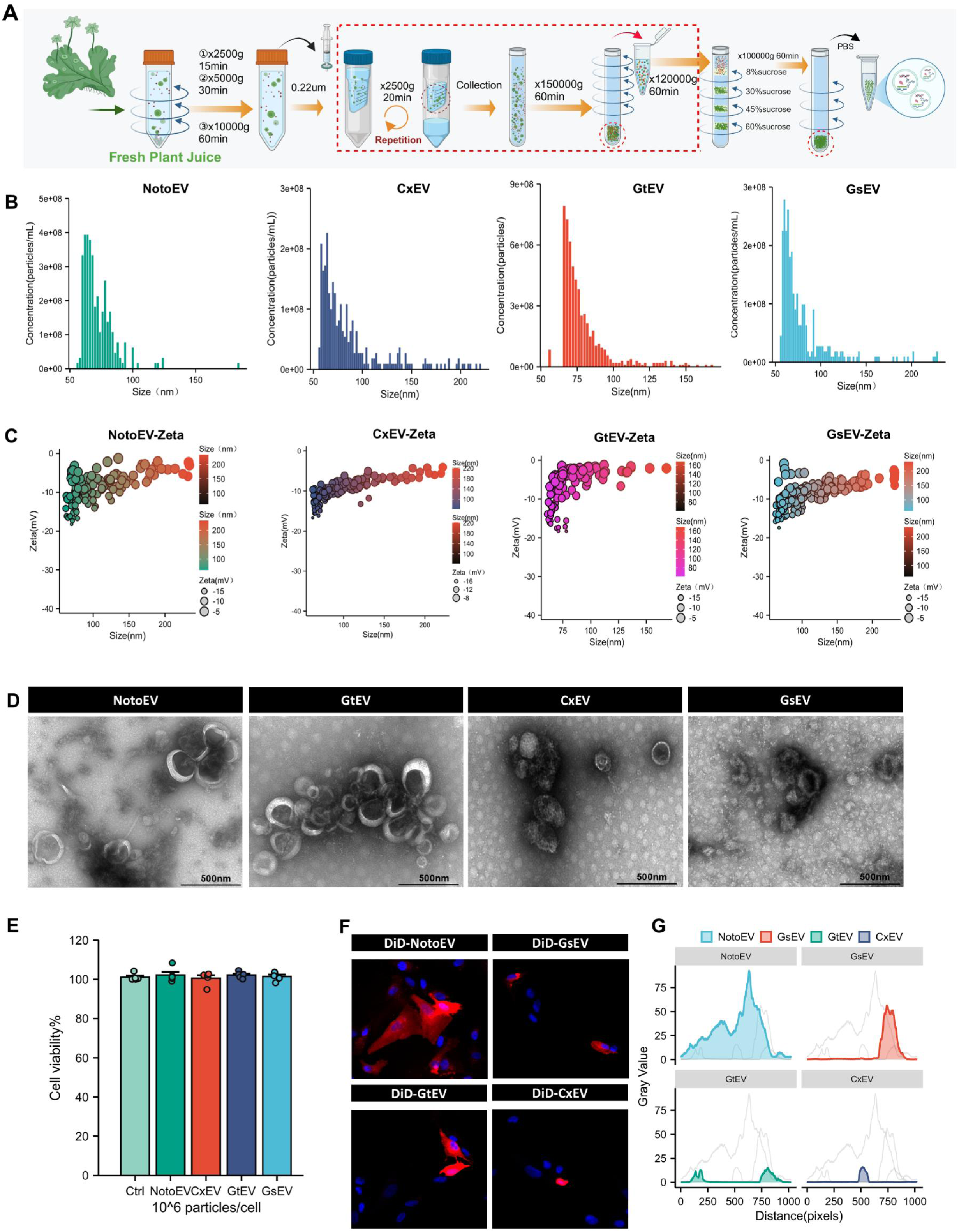
Isolation optimization, physicochemical characterization, and cellular uptake of PEVs. (A) Schematic of the optimized PEV isolation protocol, with key modifications (dashed red boxes) enhancing extraction efficiency. (B–D) Physicochemical properties of GtEVs, CxEVs, NotoEVs, and GsEVs: (B) Size distribution, (C) zeta potential, and (D) TEM images showing characteristic cup-shaped morphology. (E) Biocompatibility assessment of PEVs via SH-SY5Y cell viability assay (n = 5). (F–G) Cellular uptake evaluation: (F) Confocal microscopy images of DiD-labeled PEVs and (G) quantitative fluorescence intensity analysis.

For the preliminary characterization of the four types of plant EVs, a nano- Coulter dual-channel particle analyzer, based on the Coulter principle, was used to assess particle concentration and size. The concentrations of all four EV types reached an order of magnitude of 10^12^ particles/mL. Among them, the NotoEVs exhibited the highest concentration, at 7.57 × 10^12^ particles/mL. Moreover, the particle size distribution was uniform, with an average diameter of 94 nm and an average zeta potential of –8.80 (Figures 1B-C). Transmission electron microscopy (TEM) with negative staining analysis revealed that all four types of EVs exhibited a characteristic cup-shaped morphology, and phosphotungstic acid staining clearly delineated the boundaries of the phospholipid bilayer, confirming their intact vesicular structure (Figure 1D).

To evaluate the neuro-targeting potential of EVs, we employed the 1,1’- dioctadecyl-3,3,3ʹ,3ʹ-tetramethylindodicarbocyanine perchlorate (DiD) fluorescence labeling technique to analyze the uptake efficiency of SH- SY5Y cells. The results demonstrated that the cellular uptake rate of NotoEVs was significantly higher than that of the other three EV types (NotoEVs > GsEVs > GtEVs > CxEVs), and cell viability in all exosome-treated groups was non-significantly affected (*p* > 0.05), demonstrating their good biocompatibility (Figures 1E-F).

Verification of their anti-apoptotic effects using the OGD/R model further revealed that NotoEVs significantly upregulated the expression of the anti-apoptotic protein B-cell lymphoma-2 (Bcl2) (*p* < 0.05), inhibited the expression of the pro-apoptotic proteins BCL2 Associated X Protein (Bax) and cleaved Caspase-3 (*p* < 0.05), and effectively reduced the apoptotic rate of SH-SY5Y cells (*p* < 0.001) (Figures S1A-B, Supporting Information). Notably, although all four types of EVs exhibited an anti-apoptotic trend, the effect intensity of NotoEVs was significantly greater than that of the other groups, suggesting that their unique composition may confer enhanced neuroprotective properties.

### NotoEV promotes neuronal morphological recovery and synaptic reconstruction in the OGD/R model

The morphological integrity of neurons is the structural basis for the execution of their physiological functions and the formation of neural networks. In this study, a mouse primary neuronal OGD/R model was established to simulate cerebral ischemia–reperfusion injury (Figure 2A). NotoEV treatment significantly improved the viability of ischemic neurons (Figure 2B), alleviated neuronal morphological damage, and promoted functional recovery. Live cell dynamic monitoring demonstrated that administering NotoEVs during the reperfusion period promoted progressive neuronal morphological repair, manifested as dynamic reconstruction of dendritic extensions and synaptic structures (Figure 2C). Microtubule-associated protein 2 (MAP2) was then used to label neurons (Figure 2D).

**Figure 2.**
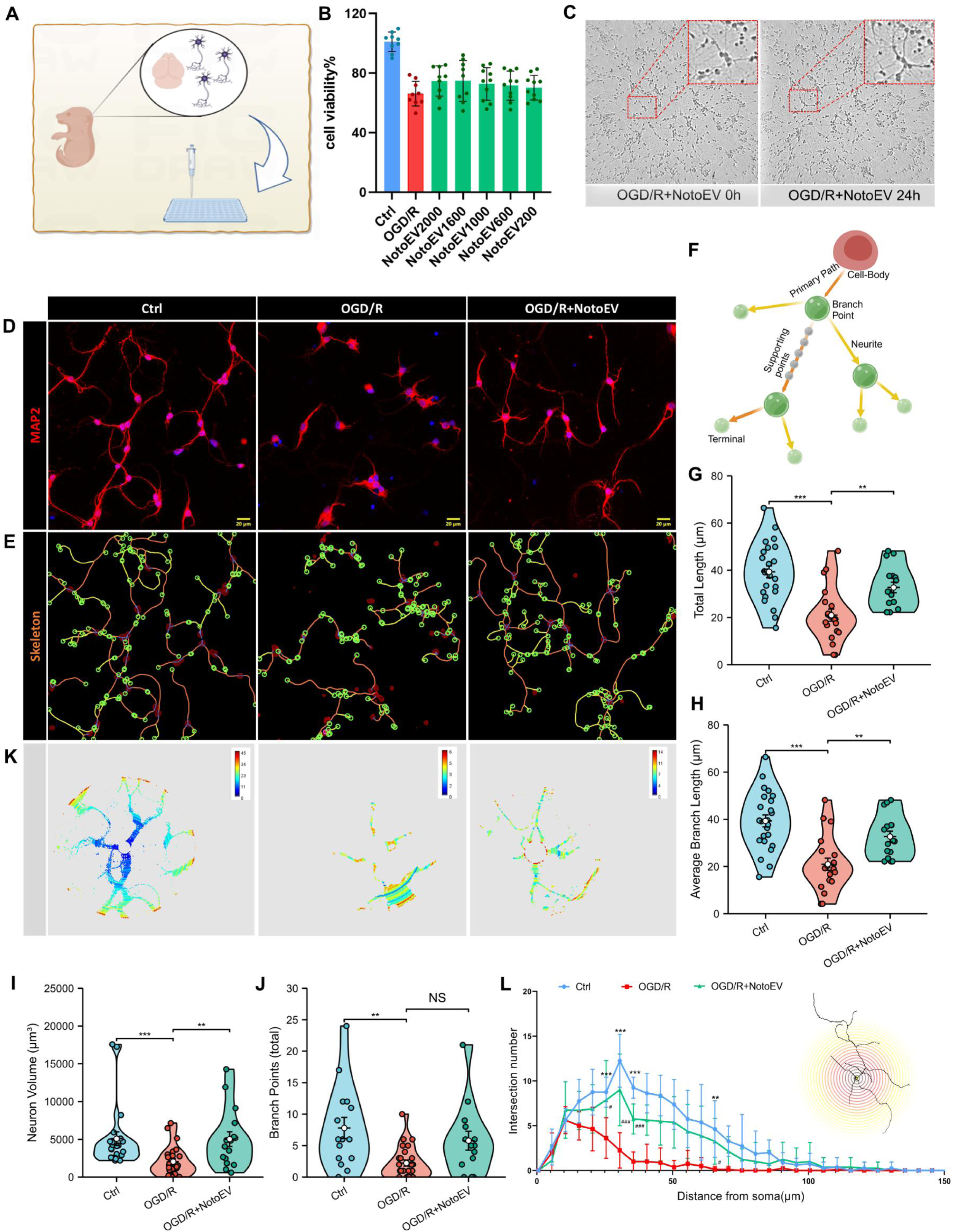
NotoEVs promote neuroprotection in primary cortical neurons by enhancing viability, neurite outgrowth, and dendritic complexity. (A) Workflow for isolation and culture of primary cortical neurons. (B) CCK-8 assay showing dose-dependent neuroprotection by NotoEVs (n = 9). (C) Time-lapse imaging of neurite regeneration during reperfusion. (D) MAP2 immunostaining demonstrating neuronal morphology post-treatment. (E – F) Arivis-based neurite reconstruction: (E) Skeletonized projections and (F) structural parameterization. (G – J) Morphometric quantifications: (G) Total neurite length, (H) average branch length, (I) neuronal volume, and (J) branch points. (K– L) Sholl analysis of dendritic arborization complexity. Data are expressed as mean ± standard error of the mean (SEM); ns: *p* > 0.05, \**p* < 0.05, \*\**p* < 0.01, and \*\*\**p* < 0.001 versus OGD/R group.

Immunofluorescence staining revealed that OGD/R treatment caused significant synaptic disruption in MAP2-positive cells, which are specific markers of neurons, and the length of dendritic branches was significantly reduced. The NotoEV treatment group effectively reversed these pathological changes, with dendritic branch length recovering to 32.70 ± 9.56 µm. Using the Arivis three-dimensional (3D) reconstruction technique for tracking and quantitative analysis of neuronal morphology, NotoEV treatment significantly improved multiple indices of neural complexity: The total dendritic length increased (*p* < 0.001), average branch length was prolonged (*p* < 0.01), neuron volume was expanded (*p* < 0.001), and the number of branch nodes increased to 75% of that in the control (Ctrl) group (Figures 2E–L). These data indicate that NotoEVs effectively repaired the neuronal morphological damage induced by OGD/R by promoting cytoskeletal reconstruction and synaptic regeneration.

### NotoEV-mediated recovery of neuronal activity in the OGD/R model

To systematically evaluate the functional repair effects of NotoEVs on neurons with ischemia-reperfusion injury, this study employed microelectrode array (MEA) technology to conduct electrophysiological dynamic monitoring of mouse cortical neurons in OGD/R and NotoEV administration groups (Figure 3A). The experimental design included three key time points: Baseline (Before), 1 h after OGD injury, and 24 h after reperfusion (After). High-throughput quantitative analysis of neuronal electrical activity was performed using high-density electrode recordings combined with the Axion Integrated Studio software analysis platform.

**Figure 3.**
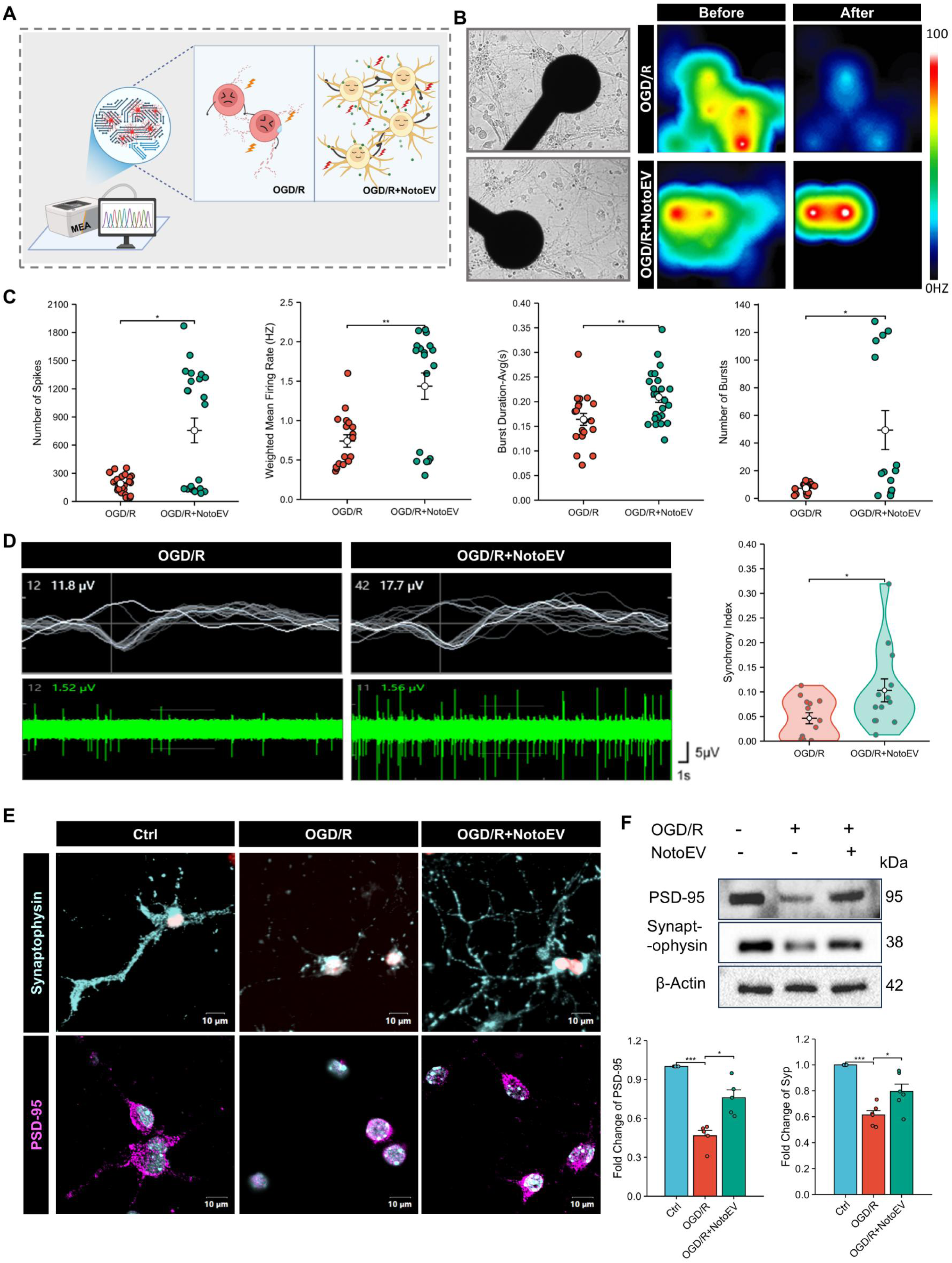
NotoEVs restore neuronal activity and synaptic integrity post-OGD/R. (A) Schematic of the multielectrode array (MEA) setup. (B – D) Neuronal network recovery: (B) Electrode-neuron interface analysis, (C) network-wide spike activity, and (D) spontaneous firing frequency. (E-F) Synaptic protein expression:(E) PSD95/synaptophysin immunofluorescence, and (F) Western blot quantification of protein levels. Data are expressed as mean ± SEM; ns: *p* > 0.05, \**p* < 0.05, \*\**p* < 0.01, and \*\*\**p* < 0.001 versus OGD/R group.

Under basal culture conditions, cortical neurons established close contact with the electrode substrate and exhibited typical radial and networked growth patterns (Figure 3B). In the baseline state, OGD/R and OGD/R + NotoEV groups maintained spontaneous discharge frequencies between 30 and 100 Hz. The number of electrophysiologically active cells in both groups decreased significantly after 45 min of OGD and 24 h of reperfusion.

Notably, the NotoEV intervention group exhibited better recovery of electrical activity; its spontaneous discharge frequency was maintained within the range of 50–100 Hz, which was significantly higher than the 30 Hz observed in the OGD/R model group (*p* <0.05). The multi-parameter analysis further revealed that NotoEV treatment significantly improved key electrophysiological characteristics of the damaged neurons: (1) The number of spontaneous discharge spikes increased; (2) the weighted average firing rate increased (*p* < 0.01); (3) the continuous discharge duration was prolonged from 0.16 ± 0.03 to 0.21 ± 0.04 s (*p* < 0.05). Notably, NotoEV intervention significantly enhanced the synchronization index of the neuronal ensemble, suggesting that it could accelerate the reconstruction of neural functions by promoting synchronized activity within the neural networks (Figures 3C-D).

Immunofluorescence and Western blot analyses revealed that OGD/R injury significantly downregulated the expression of synapsis-related proteins. Specifically, the level of the postsynaptic density marker PSD95 was reduced to 47% of that in the control group (*p* < 0.001), whereas the synaptic vesicle marker synaptophysin decreased to 61% of the control levels (*p* < 0.001). Treatment with NotoEVs markedly reversed this decline, restoring PSD95 and Synaptophysin expression to 76% and 79% of Ctrl levels, respectively (*p* < 0.05 versus OGD/R group) (Figures 3E-F). These results demonstrate that NotoEVs promote synaptic reconstruction by preserving the homeostasis of key synaptic structural proteins, thereby providing a molecular foundation for the enhancement of neuronal network electrical activity.

### NotoEV improves neuronal activity via plant miRNA-dependent regulation of stress granules (SGs) and mTOR activation

To clasify the neuroprotective mechanism of NotoEVs, we performed proteomic profiling to map their regulatory cascade on stress granule (SG) dynamics and downstream pathways (Figure 4A). NotoEVs alleviated OGD/R-induced neuronal damage, including mRNA translation arrest, mitochondrial dysfunction, and synaptic loss, via pathological stress granule (SG) disruption. Mechanistically, NotoEVs disrupted SG assembly via inhibition of G3bp2, a core scaffold protein required for liquid-liquid phase separation (LLPS), thereby preventing mRNA complex aggregation. Immunofluorescence quantification confirmed a significant reduction in G3bp2-labeled SG area (Figure 4F), validating SG disassembly as a central neuroprotective axis.

**Figure 4.**
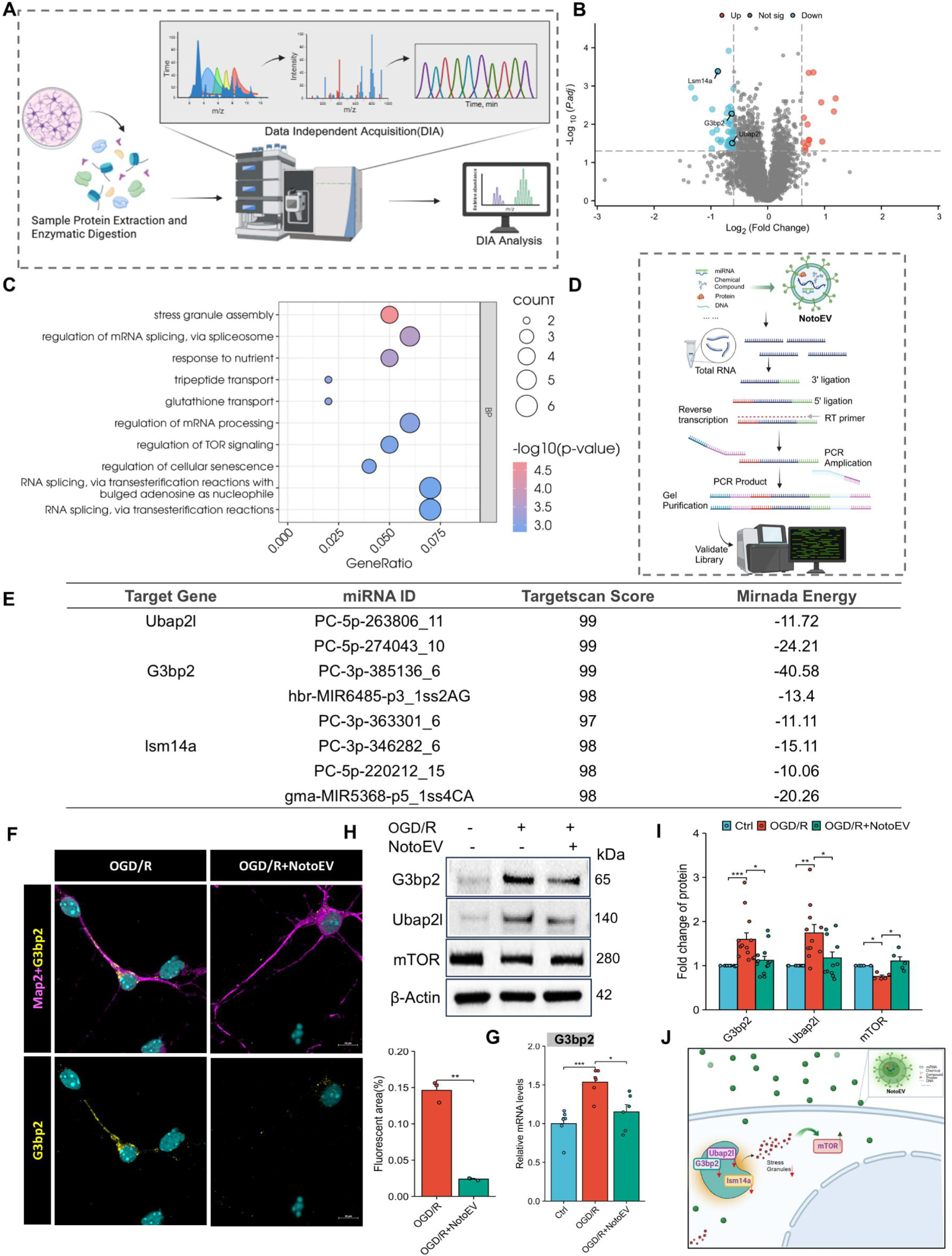
Integrated multi-omics identifies NotoEV-mediated suppression of stress granules through dual miRNA-mRNA and mTOR activation. (A) Proteomic workflow for differential protein analysis. (B) Volcano plot of significantly altered proteins (|log2FC|>1, *p* < 0.05, n=3). (C) Gene enrichment of biological processes. (D) miRNA sequencing pipeline. (E) miRNA-mRNA interaction prediction using Target Scan scores and binding energies. (F) NotoEVs reduce G3bp2 mRNA levels (qPCR). (G) Decreased G3bp2 immunofluorescence intensity Western blot and (H) corresponding quantification of stress granule associated proteins (I). (J) Proposed mechanism: NotoEVs activate mTOR to degrade G3bp2 and Ubap2l Data are expressed as mean ± SEM; ns: *p* > 0.05, \**p* < 0.05, \*\**p* < 0.01, and \*\*\**p* < 0.001 versus OGD/R group.

By regulating the Ubiquitin Associated Protein 2 Like (Ubap2l) protein, NotoEVs affect the substructure sorting function of SGs and promote the premature disassembly of immature SGs. MRNA Processing Body Assembly Factor (LSM14A), an mRNA transport protein responsible for recruiting specific mRNAs to SGs for protection, was downregulated by NotoEVs, thereby exposing key mRNAs to a stressed environment and accelerating their degradation. This downregulation also affected the assembly and maintenance of SG functions, ultimately reducing the number of SGs (Figures 4B-C).

Studies have demonstrated that [26], in addition to being a core scaffold protein of SGs, G3bp anchors the Tuberous Sclerosis Complex (TSC) complex to the surface of lysosomes and inhibits mTOR complex 1 (mTORC1) signal transduction by enhancing the GTPase activator protein (GAP) activity of Rheb GTPase. The reduction in G3bp weakens the recruitment of the TSC complex to the surface of lysosomes, relieves the inhibition of Rheb, promotes the activation of mTORC1, and increases neural activity. Our experimental results demonstrated that after NotoEV treatment, owing to the downregulation of G3bp2, the expression of mTOR protein was upregulated, which facilitated the repair of neural activity (Figures 4H-I). The data corroborate the findings from the proteomics profiling.

To analyze the active components of NotoEV, we performed deep sequencing of the miRNAs it carried and combined TargetScan (version 5.0) and miRanda (version 3.3a) for target gene prediction (screening threshold: TargetScan_score ≥ 90 and miRanda_Energy < 10) (Figures 4D-E). The results depicted that multiple miRNAs enriched in NotoEVs (including PC-5p-298830_8 and aly-miR167a-5p) targeted SG-related genes in the mouse genome through conservation of the seed region. Among these, 11 miRNAs targeted the 3’UTR region of G3bp2, 5 miRNAs regulated the post-transcriptional silencing of Ubap2L, and 7 miRNAs mediated the mRNA degradation of Lsm14a (Lsm14a) (Figure 4F). The qPCR verification demonstrated that OGD/R injury led to a 1.5-fold upregulation of G3bp2 mRNA levels in neurons (*p* < 0.001), and NotoEV intervention significantly reversed this trend by 1.2-fold (*p* < 0.05) (Figure 4G), confirming that plant miRNAs regulate the expression of target genes across species.

This study identifies NotoEVs as plant miRNA carriers that suppress G3bp2/Ubap2l-Lsm14a to disassemble pathological stress granules (SGs) and activate mTORC1 signaling, coordinating mRNA translation, metabolic recovery, and synaptic repair (Figure 4J). These findings pioneer PEV-based neuroprotection strategies targeting SG-mTORC1 crosstalk.

### NotoEV rescues mitochondrial dysfunction and neuronal apoptosis via BCL2/TOM20-mediated pathway

To systematically analyze the multi-dimensional protective effects of NotoEV in neuronal injury repair, we explored the interactive regulatory network between SGs and the mitochondrial-apoptosis axis under the stress condition of OGD/R. The study found that under OGD/R stress conditions, changes in reactive oxygen species generated by the mitochondria and the energy state affected the assembly of SGs. SGs further aggravate mitochondrial damage and promote apoptosis. NotoEVs upregulated the expression of Translocase Of Outer Mitochondrial Membrane 20 (TOM20), improved the efficiency of mitochondrial protein import, regulated the balance of Bcl2/Bax under stress conditions, promoted the recovery of mitochondrial function, reduced the release of cytochrome C, inhibited the downstream caspase cascade reaction, and affected the activation of Caspase3, thereby improving the apoptosis of nerve cells (Figures 5A–C).

**Figure 5.**
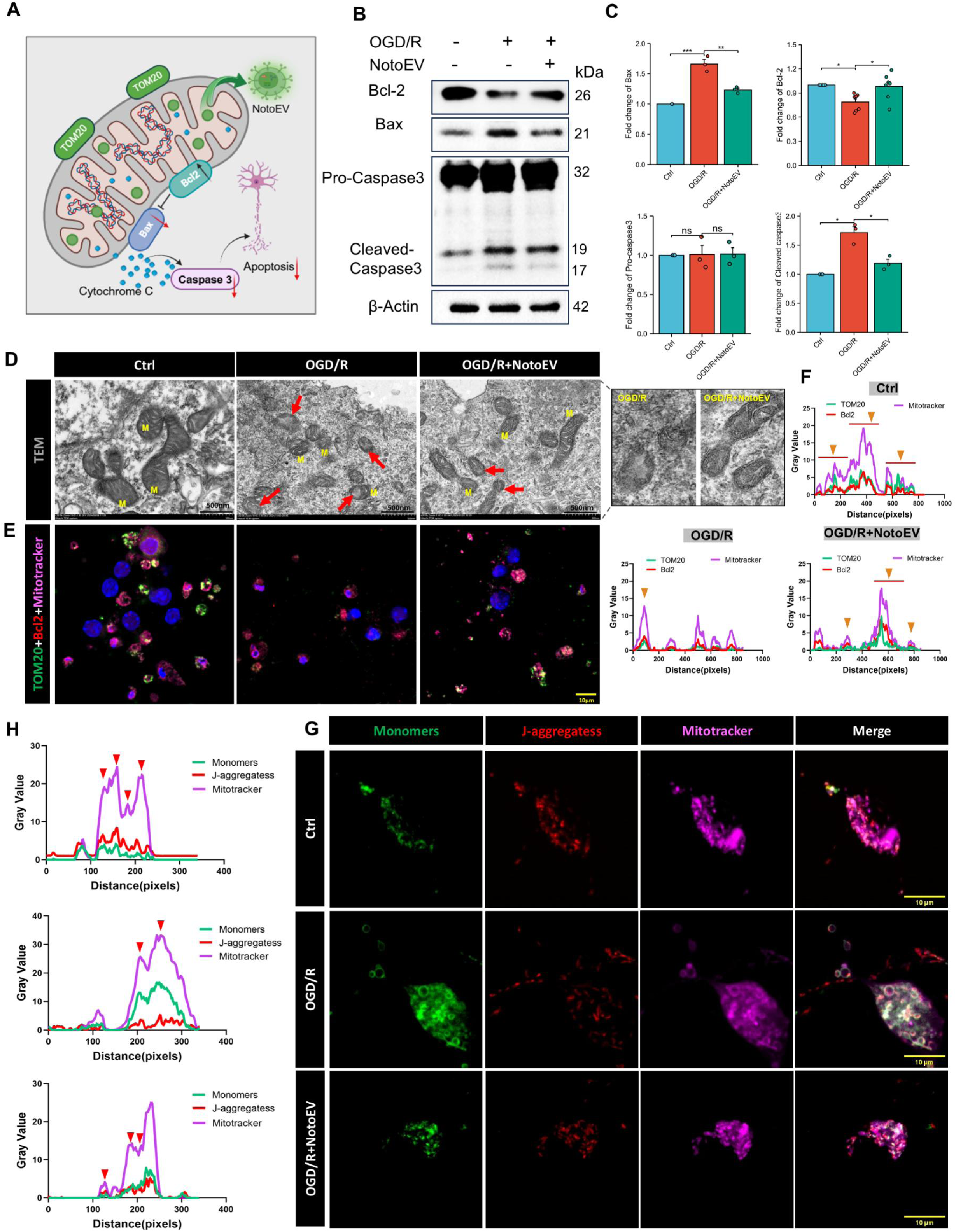
NotoEVs inhibit mitochondrial apoptosis in OGD/R-injured neurons. (A) Schematic representation of the mitochondrial protection mechanism. (B-C) Western blot and (B) corresponding quantification of apoptotic associated proteins (C). (D) TEM depicting mitochondrial ultrastructural preservation (M: mitochondria; arrows indicate crista integrity). (E-F) Bcl2 mitochondrial translocation: Mitotracker co-localization analysis. (G-H) JC-1 assay demonstrating preserved mitochondrial membrane potential. Data are expressed as mean ± SEM; ns: *p* > 0.05, \**p* < 0.05, \*\**p* < 0.01, and \*\*\**p* < 0.001 versus OGD/R group.

The results of mitochondrial TEM depicted that the mitochondria in the Ctrl group exhibited an intact overall morphology, with several inner cristae that were regularly arranged. OGD/R injury led to obvious abnormalities in mitochondrial morphology, with vacuolation, swelling, a decrease in cristae density, breakage, and disordered cristae arrangement. After administering NotoEV, the swelling state of the mitochondria was alleviated, the cristae density increased, and the breakage and disordered phenomena of the inner cristae improved (Figure 5D). Immunofluorescence co-localization analysis indicated that NotoEV significantly enhanced the co-localization of Bcl2 and TOM20, a marker protein of the outer mitochondrial membrane. Mitotracker staining demonstrated that NotoEV promoted the enrichment of Bcl2 on the outer mitochondrial membrane and directly inhibited mitochondrial outer membrane permeabilization. The increase in the fluorescence intensity of TOM20 and its co-localization with Mitotracker suggested an improvement in the efficiency of mitochondrial protein import and confirmed the recovery of nuclear-encoded mRNA translation after the reduction of SGs (Figures 5E-F).

The results of JC-1 probe detection demonstrated that, compared with the Ctrl group, OGD/R treatment significantly reduced the fluorescence intensity of red polymers and simultaneously increased the fluorescence intensity of green monomers, indicating a decrease in mitochondrial membrane potential. Fluorescence labeling with the MitoTracker mitochondrial probe further revealed that after OGD/R, the mitochondria swelled, and the neuronal mitochondria changed from a normal short rod shape to a vacuolar ring shape. Co-localization analysis demonstrated that the synchrony between JC-1 red polymer and MitoTracker fluorescence was significantly reduced, suggesting that the energy metabolism and structural integrity of the mitochondria were synchronously damaged. After treatment with NotoEVs, the fluorescence morphology of mitochondria labeled by MitoTracker was significantly improved, the fluorescence intensity of green monomers was reduced, the fluorescence intensity of red JC-1 polymers was enhanced, and synchrony with the MitoTracker signal was significantly enhanced (Figures 5G-H). NotoEVs restored mitochondrial morphology and membrane potential, resolving OGD/R-induced metabolic collapse. Concurrent apoptosis suppression and pathological stress granule (SG) clearance rescued neuronal architecture and functional recovery.

### NotoEV targets ischemic lesions and promotes neurological recovery via mTOR-dependent mitochondrial homeostasis and stress granule regulation *in vivo*

Based on previous *in vitro* experiments that confirmed that NotoEVs inhibit neuronal apoptosis by stabilizing mitochondria, we further evaluated the therapeutic potential of NotoEVs by establishing a mouse middle cerebral artery occlusion (MCAO) model. *In vivo* tracing results exhibited that DiD-labeled NotoEVs targeted brain tissues (Figure 6A). Using the PanoScanner (version 20) instrument to conduct a comprehensive scan of brain tissue slices, NotoEVs selectively accumulated in the ischemic penumbra and core infarct area (Figure 6B). Magnetic resonance imaging (MRI) and quantitative analysis indicated that compared with the MCAO model group, the infarct volume in the NotoEV treatment group was reduced by four times (*p* < 0.001) (Figures 6C and F).

**Figure 6.**
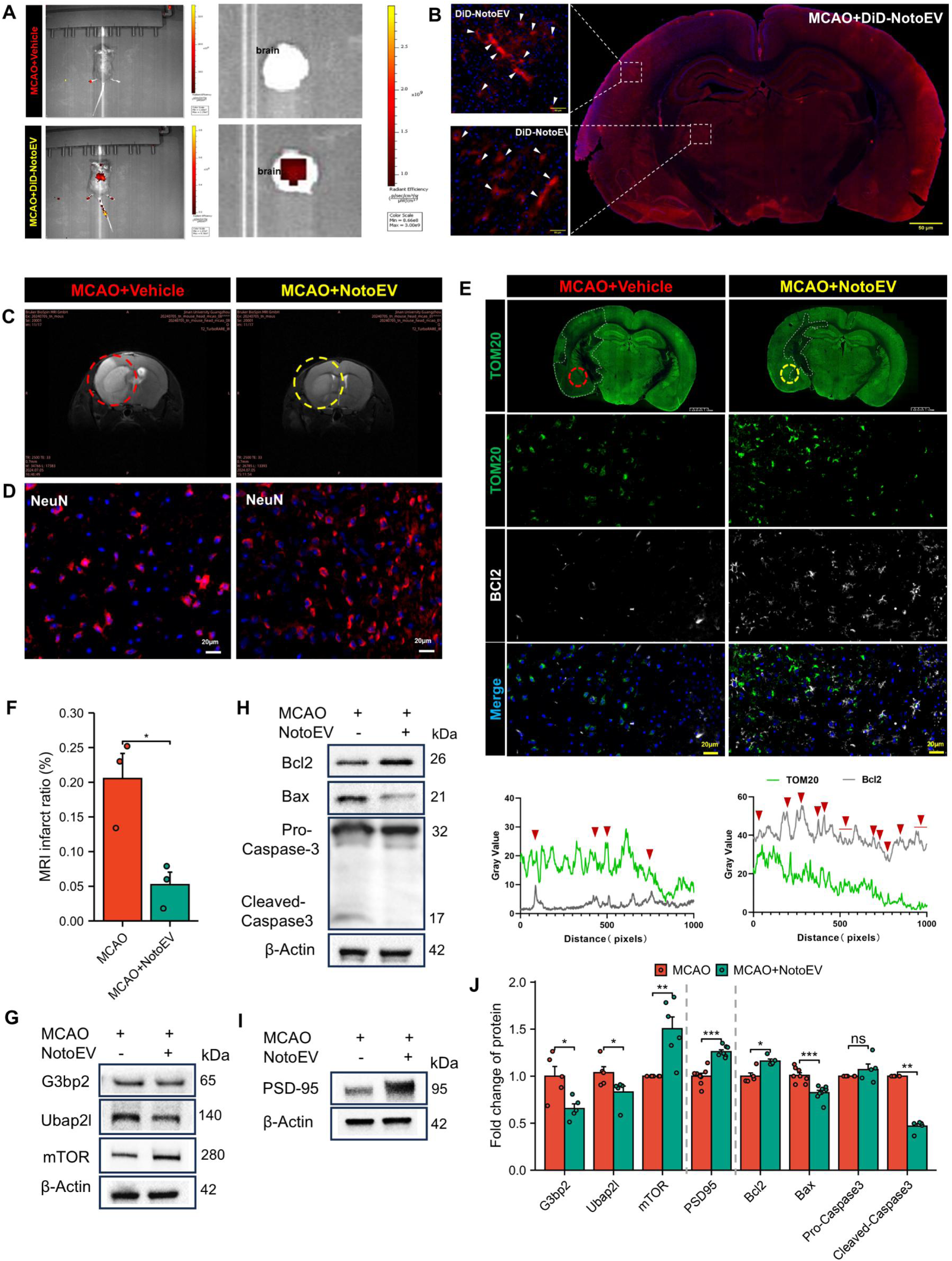
NotoEVs stimulate functional neurorestoration through mTOR activation in a mouse MCAO model. (A-B) Biodistribution of DiD-labeled NotoEVs in vivo: Whole-body imaging (A) and brain section fluorescence (B). (C) MRI quantification of infarct volume. (D-E) NeuN/Nissl staining (D) and Bcl2/TOM20 co-localization (E) in the peri-infarct zone. Boxed areas: infarct core and penumbra (white dashed line) (F–J) Molecular analyses: Western blots of SG-related (F), apoptotic (G), and synaptic proteins (H). Data are expressed as mean ± SEM; ns: *p* > 0.05, \**p* < 0.05, \*\**p* < 0.01, and \*\*\**p* < 0.001 versus MCAO group.

This structural protection was closely related to an increase in the survival rate of neurons, manifested as an increase in the number of NeuN-positive cells in the peri-infarct area (Figure 6D). Consistent with the mitochondrial strengthening effect mediated by NotoEVs in *in vitro* experiments, we mapped the cerebral infarct areas of MCAO model mice and NotoEV- administered mice according to the mitochondrial marker TOM20. Administering NotoEV significantly reduced the infarct area. The same parts of the infarct areas in the two groups were selected and magnified to demonstrate the results of the double immunofluorescence experiments with TOM20 and Bcl2. The expressions of TOM20 and the anti-apoptotic protein Bcl2 were both significantly upregulated, and co-localization analysis demonstrated that the NotoEV-administered group exhibited higher synchrony between TOM20 and Bcl2 compared with the MCAO model group (Figure 6E).

Next, we verified through Western blot that NotoEV treatment significantly downregulated the expression of the stress granule components G3bp2 and Ubap2l (Figure 6G). This result led to the enhancement of mTOR activity and promoted the upregulation of the mTOR protein. Activation of mTOR mediates multi-level neuroprotective effects. Through its influence on mitochondrial function and homeostasis, it promoted the expression of Bcl2 in the cerebral infarct area of MCAO mice, inhibited the expression of downstream apoptotic effector factors BAX and Cleaved Caspase-3 (Figure 6H), and effectively reduced neuronal apoptosis. These changes were consistent with synaptic protection, manifested as a significant upregulation of PSD95 and Synaptophysin proteins (*p* > 0.01) (Figures 6I-J). This echoed the results we observed in previous *in vitro* experiments, which demonstrated that NotoEVs enhanced neural synaptic growth and promoted the repair of neural functions.

## Discussion

PEVs are emerging as scalable biocompatible platforms for therapeutic applications that offer distinct advantages for drug delivery and regenerative medicine. Despite their promise, the low productivity and inconsistent purity of isolated PEVs have limited their application in preclinical and clinical settings. In this study, we developed an optimized isolation strategy integrating ultracentrifugation with sucrose density gradient purification, yielding PEVs at concentrations exceeding 10¹² particles/mL across four different plant species. Among these, NotoEVs exhibited the highest yield and the most stable cellular uptake, enabling detailed functional investigations.

Using an OGD/R model, a well-established *in vitro* system for ischemic injury, NotoEVs demonstrated superior neuroprotective effects compared to EVs from other plant sources. Consistent with previous studies on the neuroprotective properties of *Panax notoginseng* extracts [12], NotoEVs significantly enhanced neuronal survival by upregulating anti-apoptotic proteins and suppressing pro-apoptotic pathways. Morphological analyses revealed that NotoEV treatment preserved dendritic complexity and promoted synaptic reconstruction, which was corroborated by electrophysiological assays that demonstrated restored spontaneous firing and improved network synchronization.

A central mechanistic insight of this study lies in the regulation of SGs and the mTOR pathway by plant-derived miRNAs delivered via NotoEVs. SGs form in response to cellular stress, transiently sequestering mRNAs and proteins; however, persistent SGs contribute to neuronal dysfunction and cell death [13]. NotoEVs disrupt pathological SG assembly by downregulating the core nucleating proteins G3bp2, Ubap2l, and Lsm14a through a cross-kingdom RNA interference mechanism. This disruption alleviates mTORC1 inhibition, reactivates the protein synthesis pathways, and enhances neuronal recovery. The direct regulation of mammalian SG-related genes by plant miRNAs represents a novel therapeutic paradigm that expands the scope of cross-species RNA communication [14]. Notably, this miRNA-driven mechanism differentiates NotoEVs from mammalian EVs, which primarily exert their effects through protein cargo modulation [15].

Beyond cytoskeletal repair, NotoEVs confer mitochondrial protection, which is a crucial determinant of neuronal fate following ischemic injury. NotoEV treatment preserved the mitochondrial membrane potential, restored cristae structure, and inhibited apoptosis through the upregulation of TOM20 and Bcl2. These mitochondrial effects translated into reduced infarct volumes and enhanced neuronal survival in a mouse model of MCAO, confirming the therapeutic relevance of NotoEVs *in vivo*.

Compared to previous plant- or mammal-derived EV therapies, NotoEVs present several key advantages. Their plant origin ensures scalability and reduces immunogenic risks [16–17], thereby overcoming the production and safety challenges associated with mammalian EVs. Furthermore, the specific disruption of pathological SGs via miRNA-mediated regulation offers higher precision than traditional pharmacological SG inhibitors. These findings align with recent advances in plant EV engineering, such as the development of structural droplet drugs capable of enhancing BBB transcytosis [18]. In contrast to hydrophilic herbal monomers, these often exhibit poor BBB permeability [19–20], PEVs provide efficient brain targeting and enhanced bioavailability, highlighting their superiority in neurological applications [21–23].

Although the present study established NotoEVs as promising candidates for IS therapy, several limitations remain. Functional validation was restricted to acute ischemic models. Future studies in chronic stroke models characterized by persistent neuroinflammation and glial scar formation are needed to assess long-term efficacy. Moreover, expanding investigations into other neurological disorders, such as traumatic brain injury, Alzheimer’s and Parkinson’s diseases, where mitochondrial dysfunction and proteostasis impairment are prominent, could further establish the breadth of NotoEV’s therapeutic potential. Clinical translation requires rigorous evaluation of the pharmacokinetics, dosing, and immunogenicity of complex and human-relevant systems.

In conclusion, this study demonstrates that NotoEVs represent a novel, scalable, and mechanistically distinct platform for neuroprotection. NotoEVs offer a transformative approach for IS treatment and other neurodegenerative disorders by integrating morphological, functional, and molecular restoration through the miRNA-driven regulation of SGs and mitochondrial homeostasis. These findings lay the groundwork for future applications of PEVs as next-generation therapeutics in materials science and regenerative medicine.

## Methods

### Cell Uptake

EVs of *Panax notoginseng*, *Panax ginseng*, *Ligusticum chuanxiong*, and *Gastrodia elata* were labeled with DiD. DiD (5 μg/mL) was incubated with the plant exosome solution at 4 °C for 30 min. During this incubation period, the solution was inverted and mixed thoroughly at 5 min intervals. After incubation, the solution was divided into four ultracentrifugation tubes. Then, phosphate-buffered saline (PBS) was added to each tube to reach the appropriate volume. Each tube was then ultracentrifuged at 100,000 g for 3 h to remove free DiD dye. DiD-labeled plant EVs collected from the bottom of the tubes were resuspended and recovered using PBS. To evaluate the impact of the four plant EVs on cell viability, SY5Y cells were seeded in 96-well plates at a density of 8,000 cells per well and cultured at 37 °C under 5% CO_2_ for 24 h until the cells were fully adherent. After 24 h, the medium was replaced with a PEV-containing medium. The number of PEV administrations was set to 10^6^, 10^5^, 10^4^, 10^3^, and 10^2^ particles/cell, respectively, with 5 replicate wells prepared for each concentration. After continuous culturing for another 24 h, the CCK8 solution was added, and the absorbance at 450 nm was measured to calculate cell viability and determine the appropriate administration concentration.

To assess the uptake rates of the four plant EVs, SY5Y cells were seeded in confocal culture dishes. When the cell density reached 80%, DiD-labeled NotoEVs, GtEVs, GsEVs, and CxEVs were added to the medium at 10⁶ particles/cell. After continuous culture for 6 h, cells were fixed with 4% paraformaldehyde (PFA). An anti-fluorescence quenching agent containing DAPI was then added, and the samples were imaged using confocal microscopy.

### Microelectrode Array (MEA)

Primary neurons were inoculated with 10 µL of cell suspension at a density of 8×10^4^ cells/well onto a CytoView MEA 48-well microelectrode array test plate (Axion Company) pre-coated with poly-D-lysine/laminin. After incubation at 37 °C under 5% CO_2_ for 1h, a neurobasal medium was added to bring the total volume to 200 µL. Half of the neurobasal medium was replaced every three days. When the cells reached 8 days *in vitro*, an MEA system (Axion Biosystems) was used to collect electrical signals for 30 min to establish baseline values. The medium was then replaced with sugar-free DMEM, and 5% CO_2_ in the MEA instrument chamber was replaced with a mixed gas composed of 94% N_2_, 1% O_2_, and 5% CO_2_ for 45 min incubation. The medium was then replaced with a neurobasal medium, and NotoEVs were added at a concentration of 2,000 particles/cell. Cells were continuously cultured for 24 h during reperfusion. The untreated group served as the OGD/R group, and 7–10 wells were allocated to each experimental group. After the reperfusion period, the electrical activity of neurons in each well was recorded 24 h after OGD/R. A spike detection criterion of greater than six standard deviations above the background signals was applied to distinguish monophasic or biphasic action potential spikes from the noise. Axion Integrated Studio software was used to quantitatively process and analyze the collected data, and neuronal and neural network activities were evaluated based on parameters such as spike frequency, burst frequency, network burst frequency, and the synchrony index.

### Immunofluorescence and neuronal structure complexity

The cells were fixed with 4% PFA at room temperature for 15 min and then washed twice with PBS. The cells were then permeabilized with 0.5% Triton X-100 for 15 min and blocked with 2% bovine serum albumin for 30 min.

The cells were then incubated with primary antibodies overnight at 4 °C. After incubation, the cells were washed thrice with PBS for 5 min each and then incubated with a mixture of fluorescent secondary antibodies at room temperature for 1 h. The cells were then washed three times with PBS for 5 min each, and a mounting medium containing DAPI was added. The samples were imaged using an Olympus microscope.

The cells co-stained with G3BP and MAP2 were imaged using a Zeiss high-resolution microscope, whereas those stained with MAP2, PSD95, Synaptophysin, and Bcl2 were imaged using an Olympus microscope. For frozen brain tissue sections, after undergoing tissue permeabilization treatment for 30 min, they were directly incubated with fluorescent Nissl staining solution for 20 min and then imaged and observed.

### Live cell imaging

Neurons were seeded in 35-mm culture dishes at a density of 3.5 × 10^6^ cells/dish. The OGD model was established after 5 days of culture. During the reperfusion period, NotoEVs were administered at a dose of 2,000 particles/cell, and neurons were further cultured at 37 °C in a 5% CO_2_ incubator for 24 h. During this period, a live-cell imaging instrument was used to record the cell growth status at 10-min intervals.

### Mitochondria

Mitochondrial morphology was assessed using MitoTracker, and the mitochondrial membrane potential of primary neurons was measured using the JC-1 kit. Neurons were seeded in confocal dishes at a density of 5 × 10^5^ cells/dish. The OGD model was constructed after five days of culture. Cells were divided into three groups: Ctrl, OGD/R model, and OGD/R + NotoEV treatment. After a 24-h reperfusion period, MitoTracker dye was added at a concentration of 100 nM. After 30 min of continued culture, the cells were washed twice with the medium and then incubated with the JC-1 fluorescent probe for 20 min. The cells were washed twice with buffer and observed under a super-resolution microscope. ImageJ software was used to perform a co-localization analysis of the fluorescence signals of MitoTracker, JC-1 monomers, and polymers.

### Proteomics

Neurons from Ctrl, OGD/R, and NotoEV groups were collected, with three biological replicates per group, and total proteins were extracted. For every 5 µL of the sample, 30 µL of lysis buffer (6 M urea and 2 M thiourea) was added and mixed thoroughly. Then, 5 µL of 0.2 M TCEP and 2.5 µL of 0.8 M IAA were added to facilitate protein denaturation, reduction, and alkylation reactions. After enzymatic digestion and desalting, liquid chromatography-mass spectrometry was performed. A UHPLC system (Bruker Daltonics, Germany) was used for liquid chromatography, and a time-TOF HT mass spectrometer (Bruker Daltonics, Germany) was employed for data acquisition. FragPipe database search software was used to analyze the data.

### Electron Microscopy of Mitochondria

The neurons were collected and fixed using an electron microscope fixative at room temperature for 30 min. The fixed cells were washed with PBS and embedded in agarose. They were then fixed with 1% OsO_4_ (PBS, pH 7.4) for 2 h and sequentially dehydrated with gradient ethanol and acetone at room temperature. The cells were then embedded in Embed 812 (SPI, 90, 529-77-4) resin, and the cell sections were placed on 150-mesh copper grids using a formvar film with a thickness of 60–80 nm. Finally, the sections were stained with 2% uranyl acetate in saturated alcohol for 8 min and 2.6% lead citrate for 8 min. After drying overnight at room temperature, the samples were observed and photographed using a transmission electron microscope (Hitachi, HT7800).

### Animals

Fifty C57BL/6 mice were obtained from Zhuhai Bestest Biotechnology Co. Ltd (Zhuhai, China). The mice were housed in an environment with precisely controlled temperature and humidity and had ad libitum access to food and water. Animal group assignment followed a predetermined randomization sequence generated by statistical software. To ensure allocation concealment, the randomization schedule was implemented by a researcher not involved in subsequent experimental procedures. This study was conducted in strict accordance with the recommendations of the National Institutes of Health (NIH) Guidelines for the Care and Use of Laboratory Animals (NIH Publication No. 8023, revised in 1978). All experiments were performed in strict conformity with the guiding principles of animal experimentation at Jinan University. Efforts were made to minimize the total number of animals used and alleviate potential pain and distress as much as possible.

### MCAO reperfusion

C57BL/6 mice were anesthetized using isoflurane gas (5% for induction and 1.5% for maintenance; obtained from RWD Life Sciences, Shenzhen, China). After disinfection with iodine tincture, a median incision was made in the neck region to carefully dissect and separate the left common, internal, and external carotid arteries. The middle cerebral artery occlusion method was used to establish a focal ischemia model. Specifically, a nylon wire with a silicone tip (model 2000AAA; Jialing, Guangzhou, China) was inserted into the middle cerebral artery and maintained for 60 min to induce focal ischemia. The nylon wire was then withdrawn to allow for reperfusion. Immediately after the operation, 1 mL of normal saline was injected into the abdominal cavity. The mice were then placed in a 37 °C incubator for 15 min before returning to their cages.

### *In vivo* cellular uptake of NotoEV

Four mice were administered 30 µL of Notoginseng small extracellular vesicle suspension (with a concentration of 7.31 × 10^12^ particles/mL) via nasal drip therapy. Another four mice were administered 30 µL of PBS using the same nasal drip approach. All eight mice underwent MCAO 20 min after nasal drip administration. After undergoing an MRI scan, the hearts of the mice were then perfused with ice-cold saline, and the whole body was fixed with 4% PFA. The brains were carefully collected and placed into 15 mL centrifuge tubes. They were then dehydrated using sucrose gradients of 15%, 20%, and 30%, followed by embedding in an optimal cutting temperature compound. The embedded brains were sliced into 10 µm sections using a cryostat (Thermo Fisher Scientific, Waltham, MA, USA).

### MRI for mice

MRI scans of the mice were performed using a 9.4 T small animal MRI scanner (Bruker PharmaScan). Mice were anesthetized by inhalation of 2% isoflurane via a nose cone, and their body temperature and respiratory rate were continuously monitored during the scanning process. T2-weighted imaging (T2WI) was performed 24 h after MCAO was established. The specific scanning parameters were as follows: A two-dimensional fast-spin echo sequence with a repetition time/echo time of 3500/33 ms and an average of two repetitions, a field of view of 20 × 20 mm, 17 axial slices with a slice thickness of 1 mm, and a matrix size of 256 × 256. The scanned area, excluding the olfactory bulb, was positioned over the brain. T2WI images were scanned and quantified using the 3D Slicer software (https://www. slicer. org/) at the same scale and for the same brain slices as the MCAO mouse images. 3D images were reconstructed using a 3D slicer based on T2WI images, and the infarct and non-infarct regions were identified through threshold adjustment.

### *In vivo* tracing study of NotoEV

Mice (n=3 per group) received intranasal administration of DiD-labeled NotoEVs (1.5 × 10^12^ particles/mL in 30 μL) or PBS (30 μL) 20 min prior to transient middle cerebral artery occlusion (MCAO). Under anesthesia, mice underwent in vivo imaging using an IVIS Lumina III system (PerkinElmer) to track systemic biodistribution. Following euthanasia, ex vivo fluorescence imaging of brain sections was performed to assess trans- blood-brain barrier (BBB) penetration and parenchymal localization of NotoEVs within ischemic lesions.

## Data analysis

All statistical analyses were completed using online bioinformatics tools and R packages provided by Xiantao Academic. All statistical tests were performed in the R environment. The customized R packages of Xiantao Academic were used to perform various data statistics and data visualization plots. The software used was R (version 4.2.1), and the R packages used were ggplot2 (version 3.4.4), stats (version 4.2.1), and car (version 3.1-0) software. Appropriate statistical methods were automatically selected for statistical analysis based on the characteristics of the data format. Data that did not meet statistical requirements were not subjected to statistical analysis. The ggplot2 package was used for data visualization.

For two-group data, the t-test was used when the data met the requirements of normality and homogeneity of variance; the Welch t-test was used when the data met normality but not homogeneity of variance, and the Wilcoxon rank-sum test was used when the data did not meet normality. For three or more groups of data, one-way analysis of variance (ANOVA) was used when the data met the requirements of normal distribution and homogeneity of variance. Welch’s one-way ANOVA was used when the data met normality, but not homogeneity of variance, and the Kruskal– Wallis test was used when the data did not meet the normal distribution. Sholl analysis data were statistically analyzed using two-way repeated-measures ANOVA. All representative images were selected without bias and had typical characteristics of the data or an overall trend.

## CRediT authorship contribution statement

Hongcheng Mai and Dan Lu conceived the idea and designed the experiment. Yuanyuan Yu, Na Tan, Zhifeng Xu, performed experiments. Zhijian Tan, Tao Wang, Huimin Liu analysed the data. Yamei Tang improved the manuscript.

## Acknowledgments

This work was supported by grants from Guangdong Basic and Applied Basic Research Foundation (2025B1515020086), Sun Yat-Sen University Hundreds of Talent Program (1320324001), Fundamental Research Funds for the Central Universities, Sun Yat-sen University (24qnpy309) to Hongcheng Mai; STI 2030 Major Projects (2022ZD0211603), Guangdong S&T Program (2023B0303040003), Science and Technology Program of Guangzhou(2023A03J0708) and National Natural Science Foundation of China (82330099) to Yamei Tang; the ‘‘Guangdong Special Support Program” (referred to as the ‘‘Guangdong Tezhi Plan”) from the Provincial Health Commission (0720240214), the National Natural Science Foundation of China (82271304, 81801150, 81971121, 82171316 and 81671167) to Dan Lu.

## Ethics approval

All animal procedures were approved by the Institutional Animal Care and Use Committee of Jinan University (approval ID:20210702-16). The study protocol conforms to the ethical guidelines of the NIH Guide (NIH Publications No. 8023, revised 1978) for the Care and Use of Laboratory Animals.

## Data availability statement

The mass spectrometry proteomics data have been deposited to the ProteomeXchange Consortium (https://proteomecentral.proteomexchange.org/cgi/GetDataset?ID=PXD064044) via the iProX partner repository [24–25] with the dataset identifier PXD064044. The miRNA sequencing data of NotoEVs have been deposited in the NCBI Sequence Read Archive (SRA) under BioProject accession number PRJNA1290867.

